# Nanopore-based single molecule sequencing of the D4Z4 array responsible for facioscapulohumeral muscular dystrophy

**DOI:** 10.1101/157040

**Authors:** Satomi Mitsuhashi, So Nakagawa, Mahoko Takahashi Ueda, Tadashi Imanishi, Martin C Frith, Hiroaki Mitsuhashi

## Abstract

Subtelomeric macrosatellite repeats are difficult to sequence using conventional sequencing methods owing to the high similarity among repeat units and high GC content. Sequencing these repetitive regions is challenging, even with recent improvements in sequencing technologies. Among these repeats, a haplotype carrying a particular sequence and shortening of the D4Z4 array on human chromosome 4q35 causes one of the most prevalent forms of muscular dystrophy with autosomal-dominant inheritance, facioscapulohumeral muscular dystrophy (FSHD). Here, we applied a nanopore-based ultra-long read sequencer to sequence a BAC clone containing 13 D4Z4 repeats and flanking regions. We successfully obtained the whole D4Z4 repeat sequence, including the pathogenic gene *DUX4* in the last D4Z4 repeat. The estimated sequence accuracy of the total repeat region was 99.8% based on a comparison with the reference sequence. Errors were typically observed between purine or between pyrimidine bases. Further, we analyzed the D4Z4 sequence from publicly available ultra-long whole human genome sequencing data obtained by nanopore sequencing. This technology may be a new tool for studying D4Z4 repeats and pathomechanism of FSHD in the future and has the potential to widen our understanding of subtelomeric regions.

## Introduction

Facioscapulohumeral muscular dystrophy (FSHD) is one of the most prevalent adult-onset muscular dystrophies. The genomes of most patients with FSHD have a common feature, i.e., a contracted subtelomeric macrosatellite repeat array called D4Z4 on chromosome 4q35. The D4Z4 array consists of a highly similar 3.3-kb single repeat unit. Normally, the D4Z4 array is highly methylated and forms heterochromatin. Patients with FSHD have less than 11 D4Z4 repeats ^1–3^. In Japan, the majority of patients with FSHD have less than 7 repeats ^4^. Shortening of the D4Z4 array causes the de-repression of the flanking genes as well as *DUX4*, located in the last D4Z4 repeat. The ectopic expression of *DUX4* is toxic in muscle tissues and is thought to be a causal factor for FSHD ^5–9^. In addition to the repeat number, the haplotype of the last D4Z4 repeat is important for the development of FSHD ^1^,^2^. The telomeric flanking region of D4Z4 contains the 3' UTR of *DUX4* and is called the pLAM region. The presence of a polyadenylation signal in this region allows *DUX4* expression and disease manifestation ^10^. In contrast, individuals without polyadenylation signals do not manifest the disease ^2^.

Molecular diagnosis of FSHD is commonly made by Southern blotting of genomic DNA after restriction enzyme digestion to measure the D4Z4 array length and estimate the number of repeats. Haplotype analysis requires a different probe ^1^. Sequencing of this D4Z4 array using Sanger sequencing or short-read sequencers (up to 600 bp for Illumina and IonTorrent) is technically difficult owing to the high similarity and the high GC content of the repeats. The Oxford Nanopore Technologies MinION (Oxford, UK) is a single-molecule sequencer that can produce long reads exceeding 100 kbp ^11^. Therefore, MinION sequencing may enable the determination of pathogenicity by sequencing the complete D4Z4 array.

## Results

### Nanopore-based D4Z4 sequencing using a BAC clone

The D4Z4 array on 4q35 has EcoRI sites in its flanking region. We took advantage of this restriction enzyme to excise the full-length D4Z4 repeats with flanking sequences, for a total of 49,877 bp. Both sides of the EcoRI-digested DNA fragment had unique sequences that are not found in the D4Z4 repeats (4,823 bp on centromeric side and 865 bp on the telomeric side). We used a bacterial artificial chromosome (BAC) clone containing 13 D4Z4 repeats with flanking regions on chr4 (RP11-242C23) cloned to the backbone pBACe3.6. RP11-242C23 contained multiple EcoRI sites (Figure 1a). pBACe3.6 vector-derived DNA was digested, yielding fragments of less than 10 kb (Figure 1a). We were able to easily separate the D4Z4-containing DNA fragment (49877 bp) from vector-derived DNA by agarose gel electrophoresis and gel extraction (Figure 1b). We extracted the D4Z4 array-containing DNA and subjected it to MinION 1D sequencing (Oxford Nanopore Technologies, Oxford, UK). Base-calling was initially performed using MinKNOW ver. 1.5.12 and fastq conversion was performed using poretools to obtain 20,761 reads ^12^.

**Figure 1.**
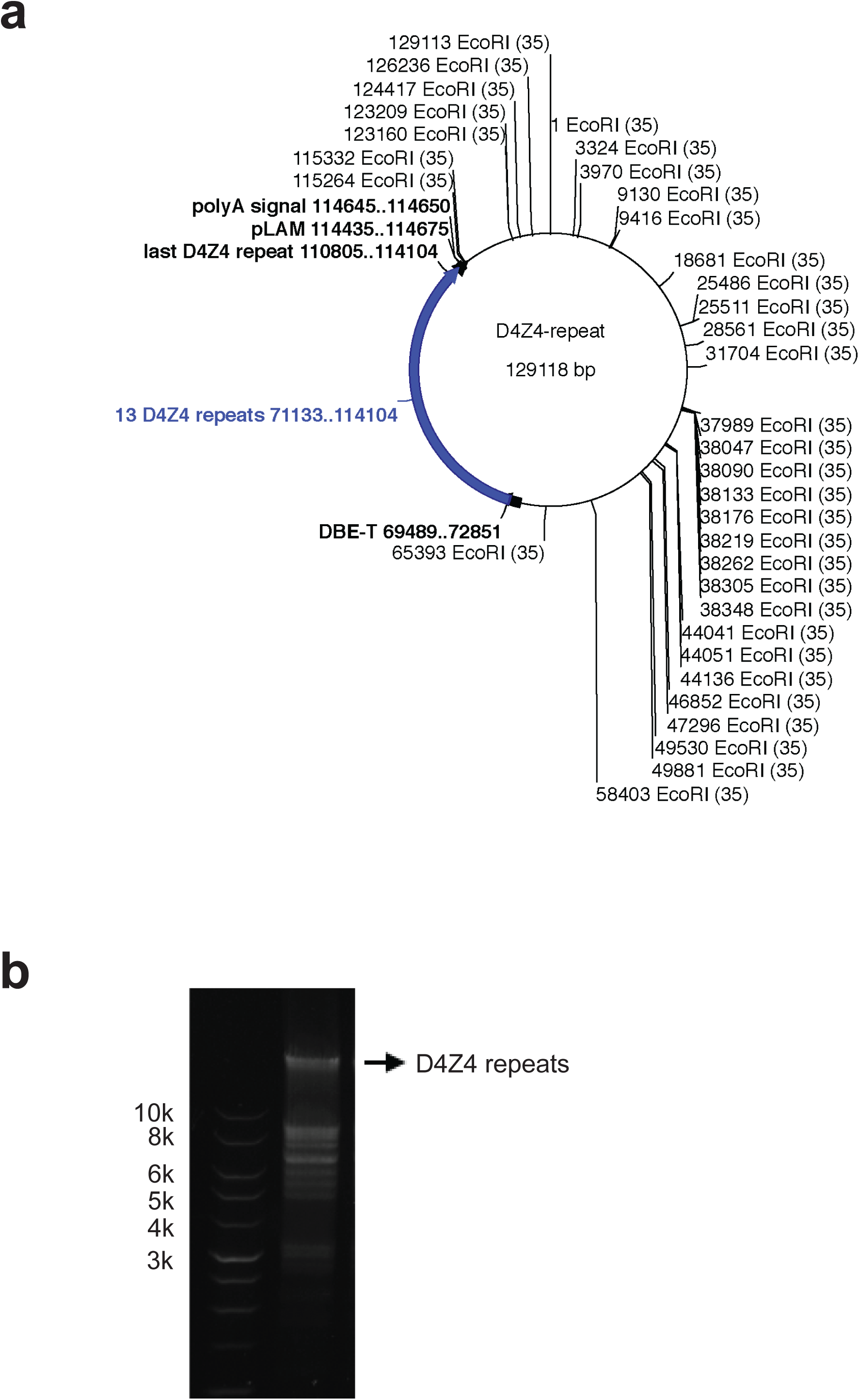
(a) Vector map of RP11-242C23 generated using Ape software. EcoRI sites are shown. The D4Z4 array with 13 repeats and flanking regions was excised using EcoRI digestion, yielding a 49877-bp product. (b) Agarose gel electrophoresis of the EcoRI-digested vector DNA. Arrow shows the band of the 49877-bp D4Z4 array.

Base-calling was not possible for 87,410 reads using the real-time MinKNOW basecaller probably due to running out of computer memory; we used Albacore (v.1.1.0) to obtain the fastq sequences in these cases. A total of 128,171 reads were obtained, with an average read length of 7,577 bp (Supplemental Table 1).

We mapped the reads to the reference BAC clone sequence (GenBank accession number CT476828.7) using LAST ^13^. Visualization of mapped reads using IGV showed coverage of the whole D4Z4 array (Figure 2). The longest read mapped to the D4Z4 repeat was 29,060 bp. The consensus sequence had an accuracy of 99.85% using last-genotype (https://github.com/mcfrith/last-genotype). The identity of the consensus sequence was 99.72% when simply employing the most common base. We also used BWA-MEM for mapping and found that the consensus sequence had a lower accuracy (99.18%). Thus, we decided to use LAST and last-genotype for subsequent analyses.

**Figure 2.**
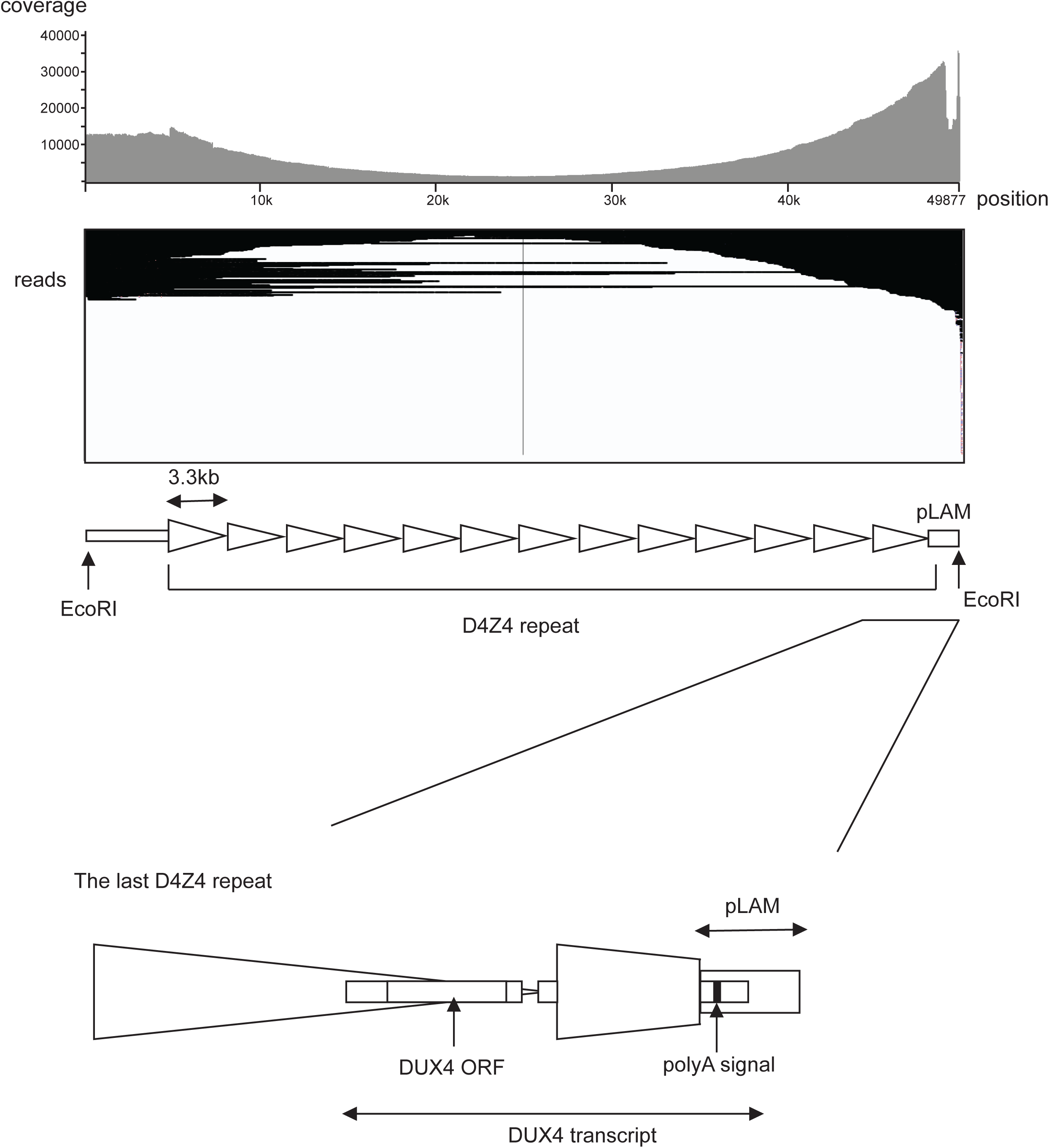
Mapped reads were visualized using IGV software. Coverage of reads is shown on the upper part of the IGV image. Scheme shows the 13 D4Z4 repeats with flanking sequences. The bottom scheme shows the enlarged last D4Z4 repeat with the pLAM region (haplotype A). This region encodes pathogenic *DUX4*. Note that IGV software has a limitation to depict reads, we showed the mapping result of 1000 randomly-chosen reads.

The haplotype of the telomeric flanking region of the last D4Z4 repeat is important for disease manifestation. There are two equally common haplotypes, A and B (Supplemental Figure 1). Haplotype A, which includes pLAM sequence, has an added polyadenylation signal at the 3' UTR of the *DUX4* gene^10^. This polyadenylation signal allows the ectopic expression of *DUX4*, which is toxic in muscle cells of patients with FSHD with the contracted D4Z4 array ^14^. Haplotype B lacks homologous sequence to pLAM. Individuals with haplotype B do not manifest the disease, despite having the contracted D4Z4 allele. Thus, it is important to identify the pLAM sequence for the molecular diagnosis of FSHD. Using MinION, we successfully sequenced the whole pLAM region with an accuracy of 100% (Figure 3). In total, 75 bases were different from the reference BAC clone sequence among the whole D4Z4 array sequence of 49,877 bp (0.15%) (Supplemental Figure 2a). Among 75 bases, 53 (70.7%) substitutions were between purines or between pyrimidines (Supplemental Figure 2b). Most of these errors were repeatedly detected at the same position in the repeats (indicated by asterisks in Supplemental Figure 2a). Interestingly, 10 out of 12 recurrent errors were seen in the CCXGG sequence at the X position. We suspect that most of these errors are likely due to base-call errors rather than random mutations in the BAC clone.

**Figure 3.**
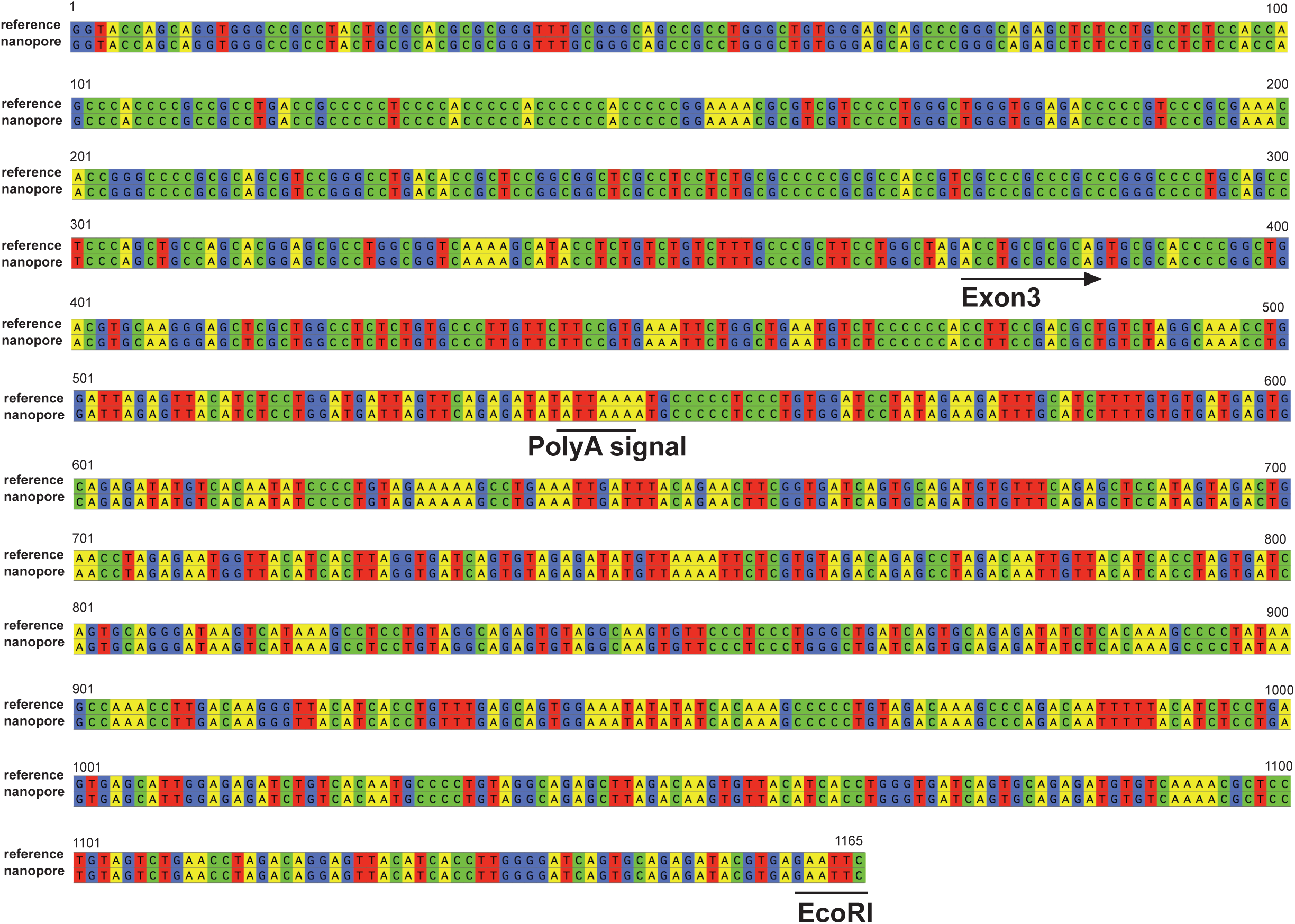
Nanopore sequence of the pLAM region. Exon 3 of *DUX4*, the 3' UTR of the gene, and polyA signal were determined with an accuracy of 100%. The upper sequence is the reference and the bottom shows the nanopore sequence.

We also compared the nanopore-sequenced *DUX4* open-reading frame (ORF) to the reference and the Sanger sequencing results for the subcloned *DUX4* ORF. The accuracy of the *DUX4* ORF sequence was 99.95% (Supplemental Figure 3).

### D4Z4 detection using nanopore-based whole human genome sequencing

We tested whether we can identify the D4Z4 array from whole genome sequencing data obtained from the MinION sequencer. We used the publicly available human reference standard genome NA12878 with R9.4 chemistry ^11^.

This project contains two sets of data. The rel3 dataset had approximately 26× coverage with an N50 length of 10.6 kb. Rel4 had 5× coverage of ultra-long reads with an N50 of 99.7 kb, indicating that rel4 contained reads that possibly cover the whole D4Z4 region. Using the ultra-long read dataset, rel4, 8 reads were aligned to the 5’ sequence of the D4Z4 repeat including p13E-11, centromeric flanking sequence (5734bp) with high confidence (mismap=1e-10, alignment length >2000 bp) using LAST. These reads were extracted and then aligned to the whole human genome (GRCh38). Two reads (read1-2) were mapped to the chromosome 10 (chr10) D4Z4 region, while two reads were mapped to the chromosome 4 (chr4) D4Z4 region (read 3-4) with high confidence: error probability <= 10^-5 (Figure4a, Supplemental Table 3). Reads that mapped to chr4 were also aligned to reference sequence for 4qA (CT476828.7), 4qAL(KQ983258.1) and 4qB(AC225782.3), and these 2 reads only aligned to the last D4Z4 of the 4qB haplotype (Supplemental Figure 4, Supplemental Figure5). To determine the number of D4Z4 repeats, those 4 reads were aligned to a single D4Z4 repeat. Both reads mapped to chr4 have 17 D4Z4 repeats and those mapped to chr10 have 20 D4Z4 repeats (Figure 4b).

**Figure 4.**
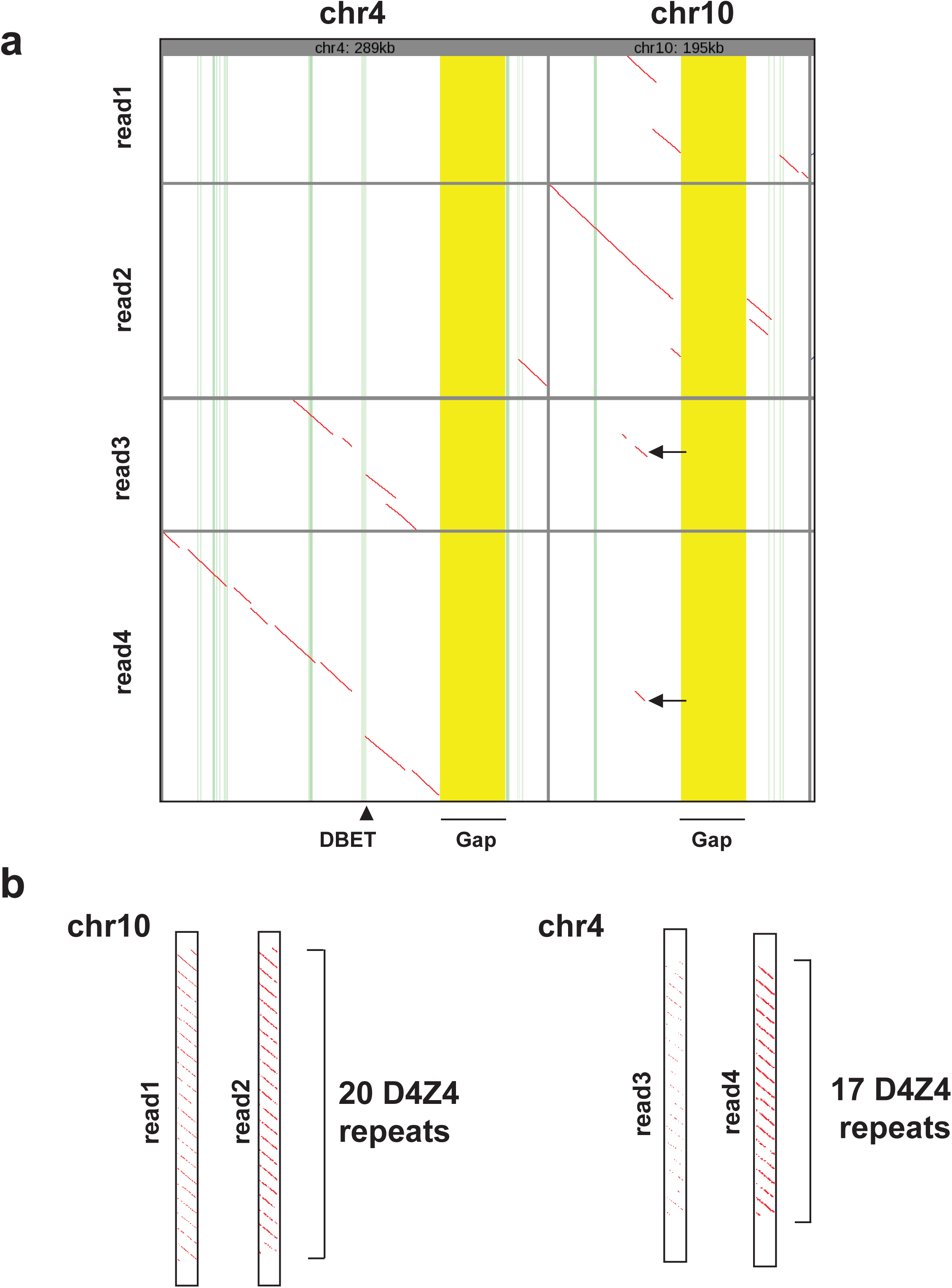
Analysis of D4Z4 repeat containing nanopore reads from whole human genome. (a) Dotplot of 4 reads that mapped to chr4 and chr10 D4Z4 and flanking regions. Gap: false gap in GRCh38 is highlighted yellow. Exons are highlighted green. DBET is highlighted light green (arrowhead). The same small portion of the chr4-read1 and chr4-read2 were aligned to chr10 (arrow). (b) Read aligned to the single D4Z4 unit using lastal and last-split revealed the repeat number. Dot plot shows 2 reads mapped to chr4 have 17 D4Z4 repeats, while those mapped to chr10 have 20 D4Z4 repeats.

## Discussion

Sequencing a highly repetitive subtelomeric region is extremely challenging. There is variation in the number of repeats among individuals and sometimes within individuals, i.e., somatic mosaicism. It has been reported that subtelomeric regions form heterochromatin, functioning as an insulator or repressor of nearby genes or preventing telomere shortening ^15^,^16^. It is important to determine the relationship between phenotypic differences and either sequence or structural variation in repeats not only to decipher the pathomechanisms of the disease, but also to obtain a deeper understanding of human genomes. Here, we applied a nanopore-based sequencer to investigate the subtelomeric repeat array associated with FSHD for the first time. In the near future, it will be feasible to search for these difficult regions to find causal relationships between the human genome and genetic diseases; even given the prevailing use of high-throughput sequencing of coding regions, the genetic causes of many diseases remain unsolved.

The disease locus of FSHD was identified at 4q35 more than 20 years ago; however, the mechanism underlying the disease has been a mystery for years and the causative genes have not been identified until recently, when accumulating evidence has shown that the misexpression of *DUX4* is associated with the disease. Further, it is still unclear whether there is any sequence polymorphism in the *DUX4* gene or flanking regions, as it is difficult to sequence the gene and the *DUX4* transcript, which is expressed at the very low levels even in the muscle tissues of patients ^14^,^17^. Since therapeutic approaches including nucleic acid drugs targeting *DUX4* mRNA are being studied ^18^,^19^, it may be useful to determine the exact *DUX4* sequence of patients for the development of effective therapies as well as an integrative diagnostic method.

Currently, the number of D4Z4 repeats is usually determined by Southern blotting using a probe that hybridizes to the centromeric flanking sequence, p13E-11 ^1^. If the patient has a deletion at this probe site, it is not possible to detect the D4Z4 repeat by Southern blotting. The Southern blotting technique is complicated and time-consuming. Alternative methods have been investigated, but are not widely used ^4^,^20^. Cost-effectiveness of the nanopore sequencers is currently uncertain. If we could enrich the D4Z4 containing DNA, one flowcell ($900 in U.S. dollars as of August 2017) is well capable of many samples to determine the repeat number and haplotype. Even a dataset with 1% of the reads of the original data in our BAC clone sequencing was enough to obtain the similar quality of consensus sequence (data not shown). MinION sequencer does not require capital investment and sequencer is provided from the company without any cost. Compared to Southern blotting, it does not require trained-skill. In addition, it produces the data in 48 hours. Although it is uncertain, considering these advantages, nanopore sequencing may be competitive with conventional techniques such as Southern blotting.

Morioka et al. sequenced D4Z4 using the PacBio sequencer ^21^ and analyzed random fragments from the BAC clone. The advantage of the nanopore sequencer over the PacBio sequencer is the ultra-long read capability^11^. It has the potential to obtain reads of more than 100 kbp, the approximate mean size of D4Z4 in healthy individuals. Currently, we could only obtain two reads that potentially cover all chr4 D4Z4 repeats from human genome data with 5× coverage using 14 flow-cells ^11^. In that paper, they were not able t o complete an alignment of the ultra-long reads using BWA-MEM. However, using LAST aligner, we could map the ultra-long reads to chr4 and chr10 and could differentiate the D4Z4 repeats from each chromosome. The advantage of the ultra-long reads is that even a single read can differentiate the highly homologous D4Z4 arrays on chr4 and chr10 because it has enough length to find the preferable alignment using the unique flanking sequence of the repeats. In addition, we could successfully align nanopore reads to a single D4Z4 repeat using the LAST aligner and last-split ^22^, which indicates the number of the repeats. This individual has 17 D4Z4 repeats with haplotype 4qB on chr4 at least on one allele, which is the normal size observed in non-FSHD individuals. As the NA12878 standard DNA originated from an individual without FSHD, this repeat number is reasonable. As FSHD patients have D4Z4 number less than 11, we think this ultra-long read sequencing may be usable to detect the disease-causing contracted D4Z4 array. If the data output for the MinION sequencer improves, it will be possible to obtain sequence data with better resolution. This approach is potentially applicable to subtelomeric regions of other chromosomes or even to centromere sequences. Interestingly, small portions of the 2 reads mapped to chr4 (read3 and read4) were also mapped to chr10. We do not know the exact reason of this (Figure 4a, arrow). One possibility is that GRCh38 does not well represent the sequence of chr4 in this region. The other possible reason is chromosomal translocation between chr4 and chr10, which is known to be frequent ^23–25^.

In nanopore-based sequencing, changes in electric current are detected as nucleotides pass through the pore. We observed that the errors tend to occur between purines or between pyrimidines, probably because they have similar chemical structures ^11^. In addition, we also observed that substitution errors tend to occur at the same nucleotide position across repeats (Supplemental Figure 2, asterisk). This may reflect the fact that the nanopore detects combinations of nucleotides and the specific combination CCXGG was prone to be misread. We anticipate further improvements of the base-calling algorithm, which will make MinION more beneficial for medical applications.

Sequencing technologies are continuously developed. During the preparation of this manuscript, the new chemistry R9.5 with the new flow-cell FLO-MIN107 was released. Considering the rapid improvements in this technique, it may not be very long before this sequencing technology is used for D4Z4 repeat analyses for patients with FSHD.

## Conclusions

Using MinION with a R9.4 flow-cell and 1D sequencing chemistry, we successfully sequenced the complete EcoRI-digested D4Z4 array from a BAC clone that contained the D4Z4 repeat region of human chromosome 4. Our deep sequencing results had an accuracy of 99.8% for the whole D4Z4 array and flanking region. This includes the pLAM region, with an accuracy of 100%, and the whole ORF of the pathogenic gene *DUX4,* with the accuracy of 99.95%, which are important regions for determining the pathogenesis. This short report may provide a basis for the future use of nanopore sequencing to deepen our understanding of highly heterogeneous subtelomeric regions that may contribute to human disease.

## Materials and Methods

### BAC clone

The RP11-242C23 human BAC clone was obtained from BAC PAC Resources Center (https://bacpaacresources.org). This BAC clone was sequenced and deposited at GenBank under accession number CT476828.7 by the Wellcome Trust Sanger Institute. It contained 13 3306-bp D4Z4 repeats.

### Preparation of D4Z4 repeats from the BAC clone

RP11-242C23 was digested using EcoRI and treated with Klenow Fragment DNA Polymerase (Takara, Shiga, Japan) at 37°C for 20 min. DNA was subjected to electrophoresis on a 0.5% agarose gel. Bands larger than the 10-kb marker (GeneRuler 1kb DNA Ladder; Thermo Fisher Scientific, Waltham, MA, USA) were excised using a razor under ultraviolet light. The DNA fragments larger than 1 kb were subjected to phenol-chloroform DNA preparation. Agarose gels were soaked in phenol and incubated for 30 min at −80°C. Then, the aqueous phase was collected and phenol-chloroform DNA preparation was performed. The EcoRI-digested whole D4Z4 repeat was enriched in the DNA sample.

### MinION 1D sequencing

Library preparation was performed using a SQK-LSK108 Sequencing Kit R9.4 version (Oxford Nanopore Technologies, Oxford, UK) using 500 ng of DNA. MinION sequencing was performed using one FLO-MIN106 (R9.4) flow cell with the MinION MK1b sequencer (Oxford Nanopore Technologies). Base-calling and fastq conversion were performed with MinKNOW ver. 1.5.12 followed by poretools or Albacore.

### Sequence alignment by LAST and BWA-MEM

Sequence reads were aligned to the EcoRI-digested D4Z4 repeat reference (Figure 1b, Supplemental material) using LAST (version 847) with the commands below ^13^,^22^,^26^;

lastdb -P8 -uNEAR -R01 rdb reference.fasta

last-train -P8 rdb reads.fasta > train.out

lastal -P8 -p train.out -m20 -j4 rdb reads.fasta | last-split > out.maf

last-genotype train.out out.maf –p 1 > genotype.txt

Sequences were also mapped to the reference genome hg19 using BWA-mem with default settings ^27^. Consensus sequences were obtained and sequence identity was calculated using UGENE ^28^. Mapped reads were visualized using IGV software ^29^.

### Subcloning of the last D4Z4 repeat

An *Escherichia coli* transformant with the RP11-242C23 human BAC clone was cultured in LB medium containing 12.5 µg/ml chloramphenicol at 37°C The human BAC clone DNA was purified using the QIAGEN Plasmid Midi Kit (Hilden, Germany) according to the “User-Developed Protocol (QP01).” Briefly, bacterial lysate from a 100-ml scale culture was passed through a QIAGEN-tip 100 column. The BAC clone DNA was eluted with buffer QF prewarmed to 65°C and concentrated by isopropanol precipitation.

To obtain the DNA clone containing the last D4Z4 repeat with the pLAM region, 50 ng of the purified BAC clone was used as a template for PCR with the forward primer 5'-cgcgtccgtccgtgaaattcc-3' and the reverse primer 5'-caggggatattgtgacatatctctgcac-3'. PCR was performed with PrimeSTAR GXL DNA Polymerase (Takara) with the following cycling conditions: 98°C for 2 min and 30 cycles of 98°C for 10 s, 60°C for 15 s, and 68°C for 30 min. PCR products were gel-purified and cloned into a pCR blunt vector (ThermoFisher Scientific) with the Mighty Mix DNA Ligation Kit (Takara). The sequence of the resulting plasmid was confirmed by Sanger sequencing with M13 forward and M13 reverse primers.

### D4Z4 sequence analysis using the ultra-long human whole genome sequence

The human whole genome sequenced by MinION sequencers was downloaded (https://github.com/nanopore-wgs-consortium/NA12878) ^11^. Using the ultra-long read dataset, rel4, the 5’ sequence of the D4Z4 repeat (5734bp) was aligned and 8 reads that have alignment with high confidence (mismap=1e-10, alignment length >2000) were extracted and then aligned to the whole human genome (GRCh38) like this ^22^,^26^;

lastdb -P8 -uNEAR -R01 db GRCh38.fasta

last-train -P8 db reads.fasta > train.out

lastal -P8 -p train.out -m50 -D1e9 hdb reads.fasta | last-split -m1e-5 | last-postmask > alns.maf

Extracted reads that mapped to chr4 were also aligned to reference sequence for 4qA (CT476828.7), 4qAL(KQ983258.1) and 4qB(AC225782.3) to determine the haplotype. To determine the number of D4Z4 repeats, those 4 reads were aligned to the single D4Z4 repeat. Alignment was visualized using last-dotplot (http://last.cbrc.jp/doc/last-dotplot.html).

## Author Contributions

SM and HM designed the study and collected experimental materials.

SM, SN, MTU, HM, MCF and TI analyzed and interpreted the data. SM drafted the original manuscript.

## Competing interests

The authors report no disclosures relevant to the manuscript.

## Web Resources

LAST: http://last.cbrc.jp

BWA: http://bio-bwa.sourceforge.net

Ape: http://biologylabs.utah.edu/jorgensen/wayned/ape/

UGENE: http://ugene.net

last-genotype: https://github.com/mcfrith/last-genotype

last-dotplot: http://last.cbrc.jp/doc/last-dotplot.html

The URL for the human whole genome sequence data for NA12878 used in this study is as follows: https://github.com/nanopore-wgs-consortium/NA12878

## Funding

This study was supported by MEXT-Supported Program for the Strategic Research Foundation at Private Universities (to SN and MTU). This work was supported by JSPS KAKENHI Grant Number JP15K19477 (to HM) and JP26700030 (to MCF).

